# Striatal fiber photometry reflects primarily non-somatic activity

**DOI:** 10.1101/2021.01.20.427525

**Authors:** Alex A. Legaria, Ben Yang, Biafra Ahanonu, Julia A. Licholai, Jones G. Parker, Alexxai V. Kravitz

## Abstract

Calcium recording via fiber photometry is commonly used as a proxy for recording population neuronal activity *in vivo*, yet the biological source of the photometry signal remains unclear. Here, using simultaneous *in vivo* extracellular electrophysiology and fiber photometry in the striatum, along with endoscopic 1-photon and 2-photon calcium imaging, we determined that the striatal fiber photometry signal reflects primarily non-somatic, and not somatic, changes in calcium.

## Main Text

Fiber photometry enables recording of bulk calcium fluctuations in genetically defined neuronal populations. While use of this technique has grown in popularity in recent years^1^, it remains unclear whether the photometry signal reflects changes in action potential firing (*i*.*e*., ‘spiking’) or non-spiking related changes in calcium. In microscope-based calcium imaging, the potential for neuropil to contaminate somatic calcium signals motivated both optical and analytical approaches for isolating somatic calcium signals from neuropil^2–5^. However, these approaches cannot be applied to fiber photometry as the technique does not retain spatial information of the emitted fluorescence.

In the striatum, fiber photometry recordings exhibit different temporal profiles than spiking when aligned to the same behavioral events^6^. Event-related changes in spiking occur in close proximity (<1 s) to actions, whereas fiber photometry signals can ramp up for several seconds before actions^6^. To investigate this discrepancy, we performed simultaneous fiber photometry and electrophysiological recordings in freely moving mice (n = 6). We expressed GCaMP6s in the striatum using either a non-specific viral strategy in wildtype mice (n = 5) or a Cre-dependent strategy in Drd1a-Cre mice (n = 1). In the same surgery, we implanted an array of 32 tungsten microwires surrounding an optical fiber for collecting both photometry signals and spiking activity (**Fig. 1a, b**). We compared simultaneously recorded photometry and multi-unit spiking (n = 73 units from 6 mice) around movement initiation (**Fig 1c; Extended Data Fig. 1**). As previously reported^6^, the photometry signal “ramped” up for ∼10 seconds before the movement, while average striatal spiking increased transiently around the start of movement (**Fig. 1c**). We further sorted units into those that were positively (n = 29) or negatively (n = 12) modulated around the movement start and observed little evidence for ramping in either population (**Extended Data Fig. 1**).

**Figure 1.**
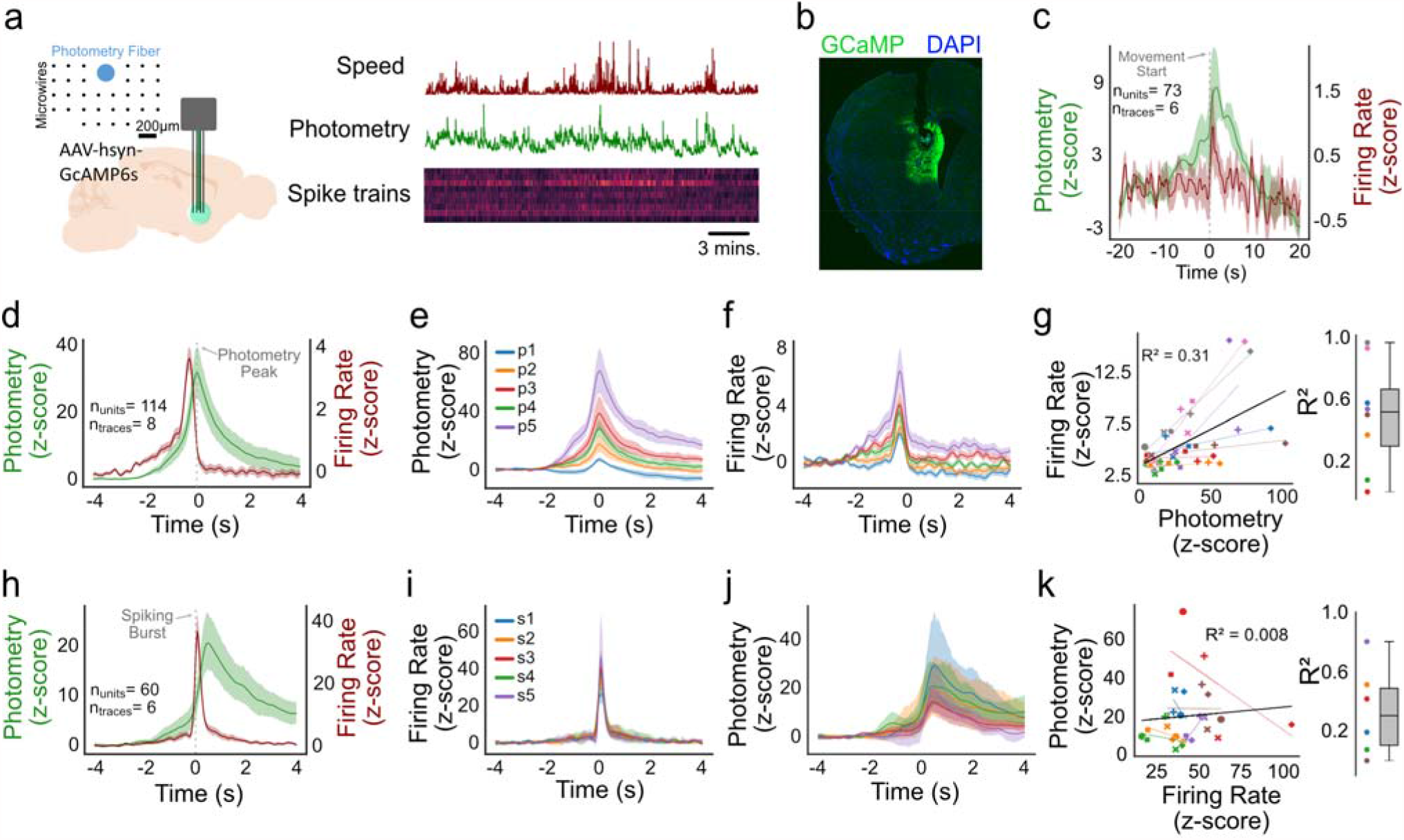
Electrophysiological correlates of fiber photometry. **a**, (*left*) Experimental setup: An optical fiber coupled to a 32-channel array (lower right) was implanted in the striatum to simultaneously record fiber photometry signals and multiunit spiking. Inset: Geometry of the array/fiber bundle. (*right*) Representative example of the three types of collected data. **b**, Photograph showing electrode damage and GCaMP expression. **c**, Average photometry and spiking activity aligned to the onset of movement. **d**, Average photometry and spiking aligned to photometry peaks. **e**, Average photometry transients split into quintiles of increasing peak prominence (p1-p5 represent photometry transients of increasing amplitudes). **f**, Spiking changes at each quintile of photometry peak prominence. **g**, (*left*) Correlation between maximum photometry and maximum spiking for each photometry peak quintile (subjects represented by color, quintiles by shape) (*right*) Quantification of (*left*). **h**, Average photometry and spiking aligned to spiking burst (s1-s5 reflect busts of increasing number of spikes). **i**, Spiking bursts of increasing amplitude. **j**, Photometry response to increasing amplitude of bursts. **k**, (*left*) Correlation between maximum photometry and spiking for burst quintiles (subjects represented by color, quintiles by shape). (*right*) Quantification of (*left*).

**Extended Data Figure 1.**
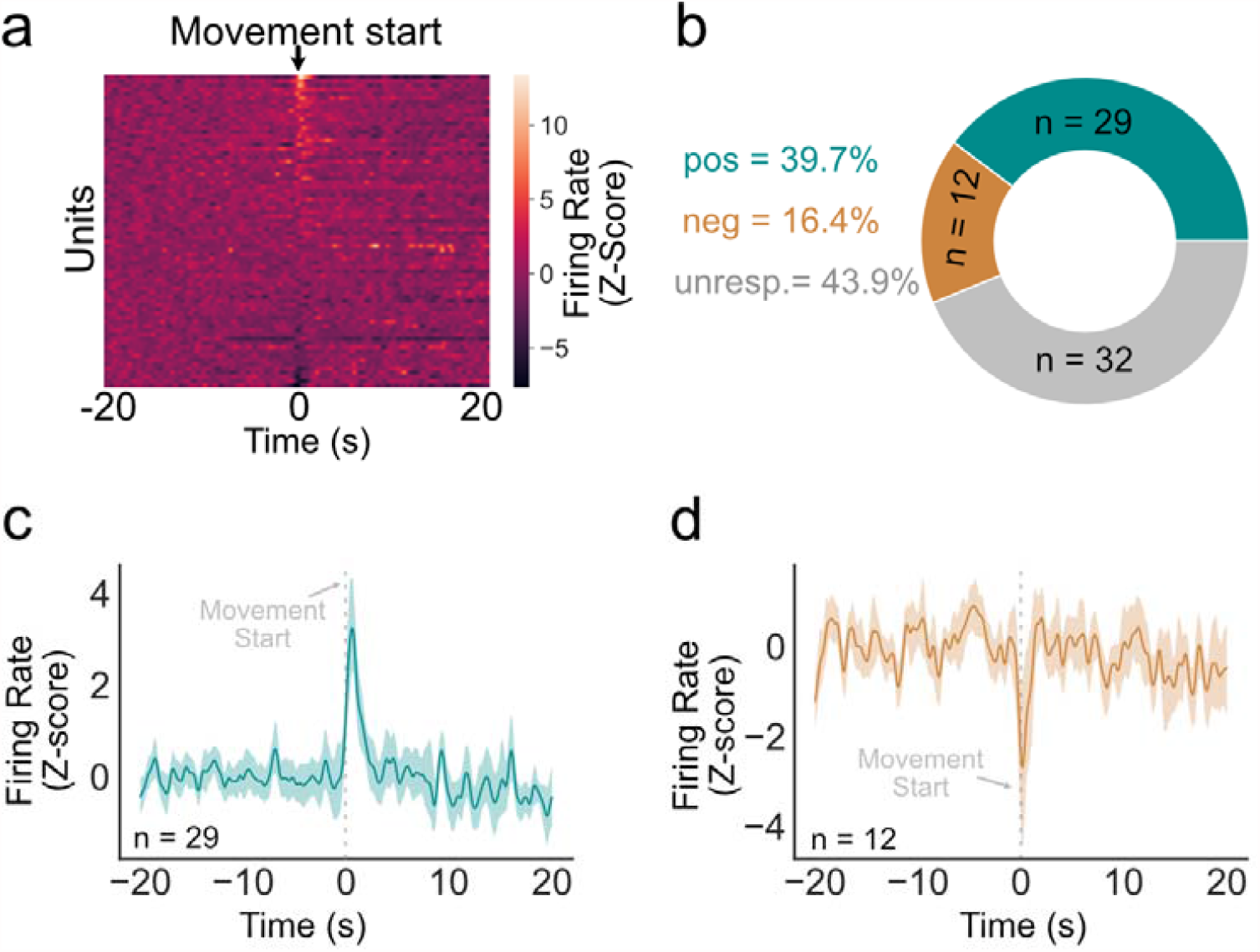
Response heterogeneity to initiation of movement. **a**, Mean spiking activity of the 73 recorded units around the start of movement. **b**, Ring plot showing the proportion of positively, negatively modulated, or unmodulated units. **c**, Average response of positively modulated units. **d**, Average response of negatively modulated units.

To further quantify the relationship between photometry and simultaneously recorded spiking, we examined changes in spiking aligned to rapid increases in fluorescence (termed: photometry transients) in 114 units recorded from 8 mice (6 WT and 2 Drd1a-Cre). Spiking activity increased during the rising phase of the photometry transients, preceding the peak of the photometry transient by ∼500ms (**Fig. 1d**), similar to prior reports^7^. We sorted all photometry transients into quintiles based on peak amplitude (p1-p5, **Fig. 1e**) and found an orderly relationship such that across all animals, larger photometry transients were associated with greater increases in spiking (**Fig. 1f**). However, this relationship was highly variable among individual animals, with the size of photometry transient explaining 32% of the overall variance in spiking activity but ranging from 0-98% across mice (**Fig. 1g**). The strength of this correlation was not dependent on number of units in each recording (**Extended Data Fig. 2a**). Moreover, there were no significant correlations between the transient amplitude (peak prominence) and absolute firing rates of recorded units, the number of spikes in bursts coincident with these transients, nor the number of units that were positively modulated coincident with photometry peaks of different sizes (**Extended Data Fig. 2b–d**). We also quantified the converse relationship, to determine whether larger spiking bursts resulted in larger changes in photometry fluorescence. Here, we observed an increase in photometry signal following spiking bursts (**Fig. 1h**) but found no quantitative relationship between the strength of bursts and the amplitude of coincident photometry transients (**Fig. 1i–k**). We conclude that photometry transients are associated with transient increases in spiking, but larger photometry transients do not necessarily reflect larger changes in spiking of the underlying population.

**Extended Data Figure 2.**
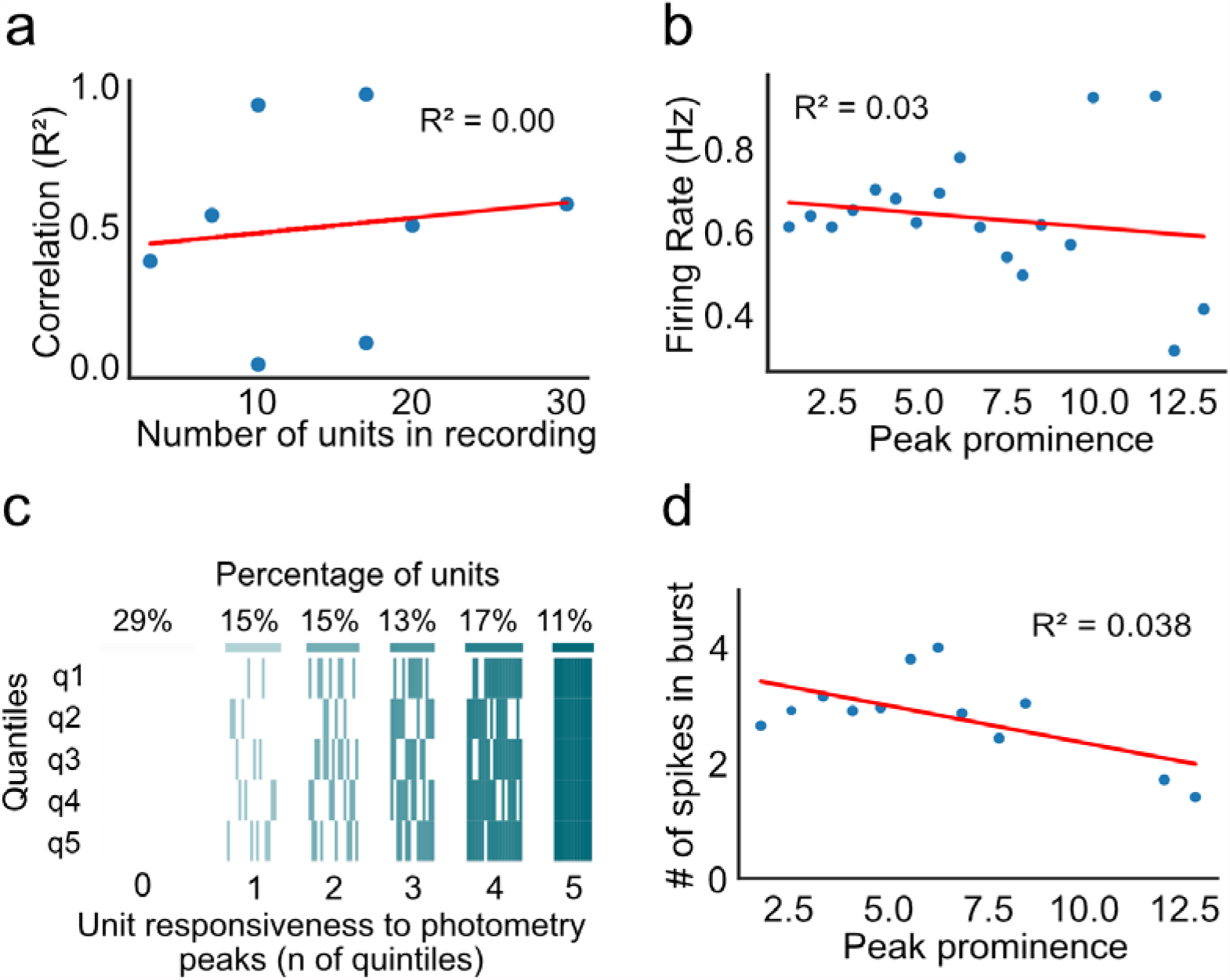
Size of photometry transient is not correlated with neuronal bursting. **a**, Correlation between photometry transient size and spiking burst strength, around bursts where there was a photometry transient present. **b**, Correlation between photometry transient size and absolute firing rate, time-locked to photometry transients. **c**, Heatmap showing number of responsive neurons to photometry peaks in each quintile. Each column represents one unit and rows represent whether that unit was responsive during each photometry quintile. Numbers on the bottom show how many quintiles each unit was responsive to (from 0-5). **d**, Correlation between the number of cells in the recording and the R^2^ value of the spiking and photometry transient relationship.

There are multiple explanations for why a larger photometry response might not reflect a larger change in spiking. For instance, ensembles of co-activated striatal neurons are spatially clustered^8,9^, so the size of observed transients may depend on the distance between the photometry fiber and an activated neuronal cluster, and not the amount of spiking in that cluster. Alternatively, the photometry signal may reflect calcium changes in the neuropil, which exhibit different spatial and temporal dynamics than somatic calcium^10^. To test this possibility, we performed microendoscopic recordings and compared the full field fluorescence signal to both somatic and neuropil signals extracted from the same imaging movies. We expressed GCaMP6m in dorsal striatum direct or indirect pathway medium spiny neurons (MSNs) of Drd1a-Cre (n = 6) or A2a-Cre (n = 6) mice, using a Cre-LoxP viral strategy (**Fig. 2a**) and recorded calcium activity from these populations through an optical guide tube containing a 1-mm diameter microendoscope gradient index (GRIN) lens^9^. We spatially cropped each of these recordings to the approximate area of a 200-µM photometry fiber for analysis, resulting in 48 movies for analysis.

**Figure 2.**
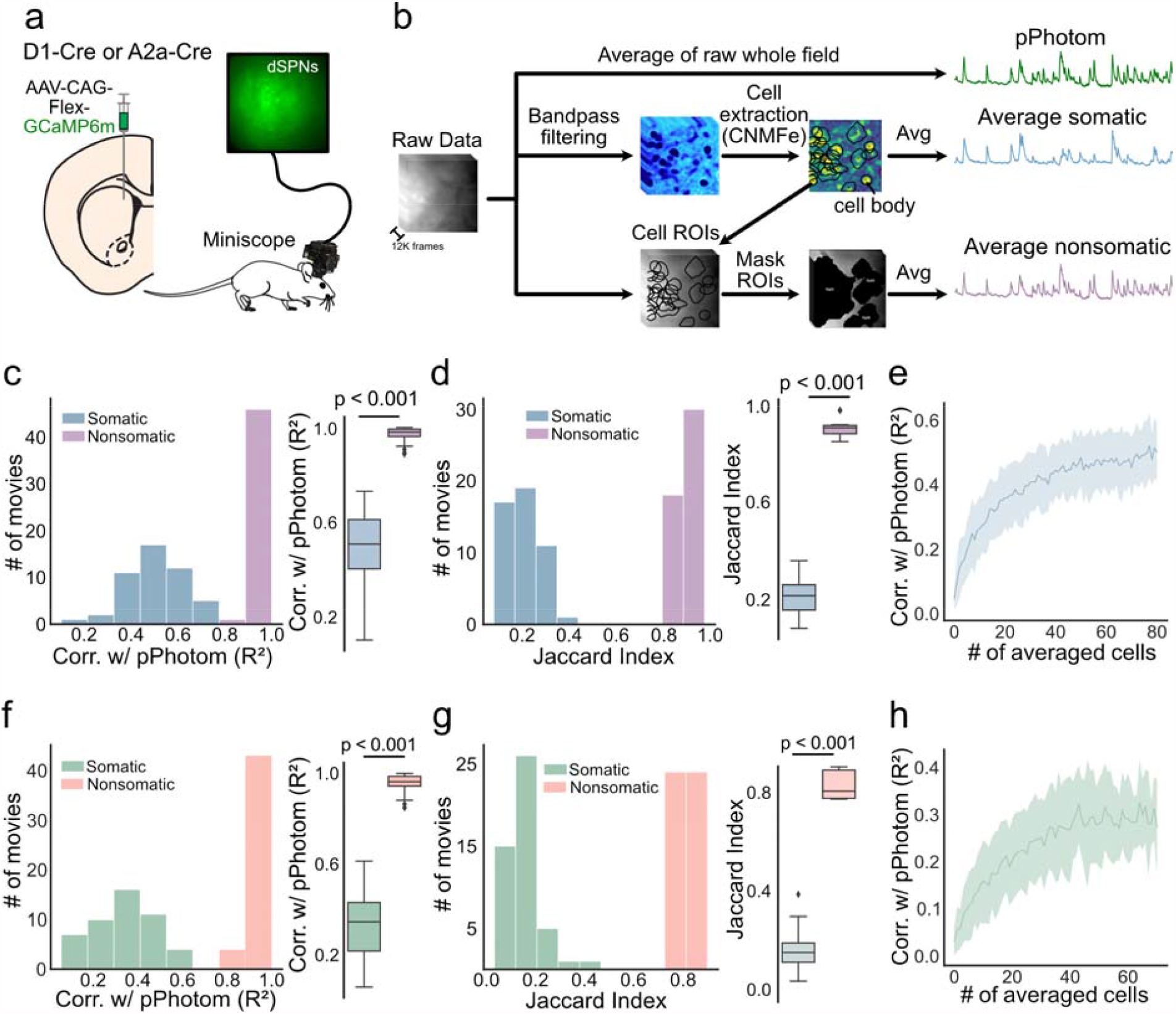
Photometry signal correlates strongly with non-somatic but not somatic changes in calcium. **a**, Experimental setup: Cre-dependent GCaMP6m was injected in the dorsomedial striatum (DMS) of D1-Cre or A2a-Cre mice and a miniature microscope was used to record neural activity. **b**, Three signals were extracted from calcium images: 1) average of the entire field (pPhotom), 2) average somatic signal through (via CNMFe cell extraction), and 3) average non-somatic signal, by masking regions were somas were present. **c**, Linear correlations between photometry and somatic and non-somatic signals in D1-positive neurons. **d**, Jaccard similarity between signal transients in the average signal, and somatic and non-somatic signals transients in D1-positive neurons. **e**, Correlation between photometry and somatic signals, as the activity of different numbers of cells was averaged. **f–h**, Same as **c–e** for A2A-positive neurons.

Three component signals were extracted from each movie (**Fig. 2b)**: 1) A proxy for the photometry signal (the average fluorescence of the raw recording, termed: pPhotom), 2) average somatic activity (individual somatic activity traces extracted with CNMFe via the CaImAn Python library^11^), and 3) non-somatic activity (regions of interest in each movie that contained no somatic activity). In both Drd1a-Cre and A2a-Cre animals, the pPhotom signal correlated more strongly with non-somatic than somatic signals (Pearson’s coefficient of determination, p < 0.001; **Fig. 2c, f**). We also calculated the Jaccard Similarity Index to compare binarized signals, independent of amplitude of changes. Again, the non-somatic signal had higher similarity to the pPhotom signal than the somatic signal (p < 0.001; **Fig. 2d, g**). To test whether the relationship between the pPhotom and somatic signals was dependent on the number of recorded somatic signals, we randomly selected increasing numbers of somatic signals (from 1-80). The correlation with pPhotom increased with additional somatic signals but plateaued at an R^2^ ∼0.5 in D1-MSNs and R^2^ ∼0.3 in A2a-MSNs (**Fig. 2e, h**), whereas the correlation with non-somatic signals averaged >0.9 for both. Based on significantly stronger correlations between pPhotom and non-somatic vs. somatic signals in all analyses, we conclude that the striatal photometry signal reflects primarily non-somatic changes in calcium.

We further hypothesized that if photometry reflects primarily neuropil, the pPhotom signal should be correlated across the whole imaging field, as dendritic and axonal arbors of medium spiny neurons can extend >500µm from somas, which are themselves only ∼10-20µm in diameter^11,12^. To test this idea, we divided the unprocessed movies into 6×6-pixel square regions (∼12 microns, similar in size to an average soma) and calculated correlations between each small region across the movies. High temporal correlations were observed between these regions in each movie (Avg. R^2^ = 0.95 ± 0.04; **Extended Data Fig. 3c, d**). In other words, regardless of where a 6×6-pixel region was located, it had a strong temporal correlation with any other 6×6-pixel region from the same movie. In addition, all individual 6×6-pixel regions correlated strongly with the pPhotom signal, independent of their location in the imaging field (Avg. R^2^ = 0.97 ± 0.02; **Extended Data Fig. 3e, f**). In contrast, individual somatic signals exhibited low temporal correlations with one another (Avg. R^2^ = 0.06±0.13), and with the pPhotom signal (Avg. R^2^ = 0.21±0.12; **Extended Data Fig. 3d–f**). These analyses further support our conclusion that photometry primarily reflects changes in neuropil calcium.

**Extended Data Figure 3.**
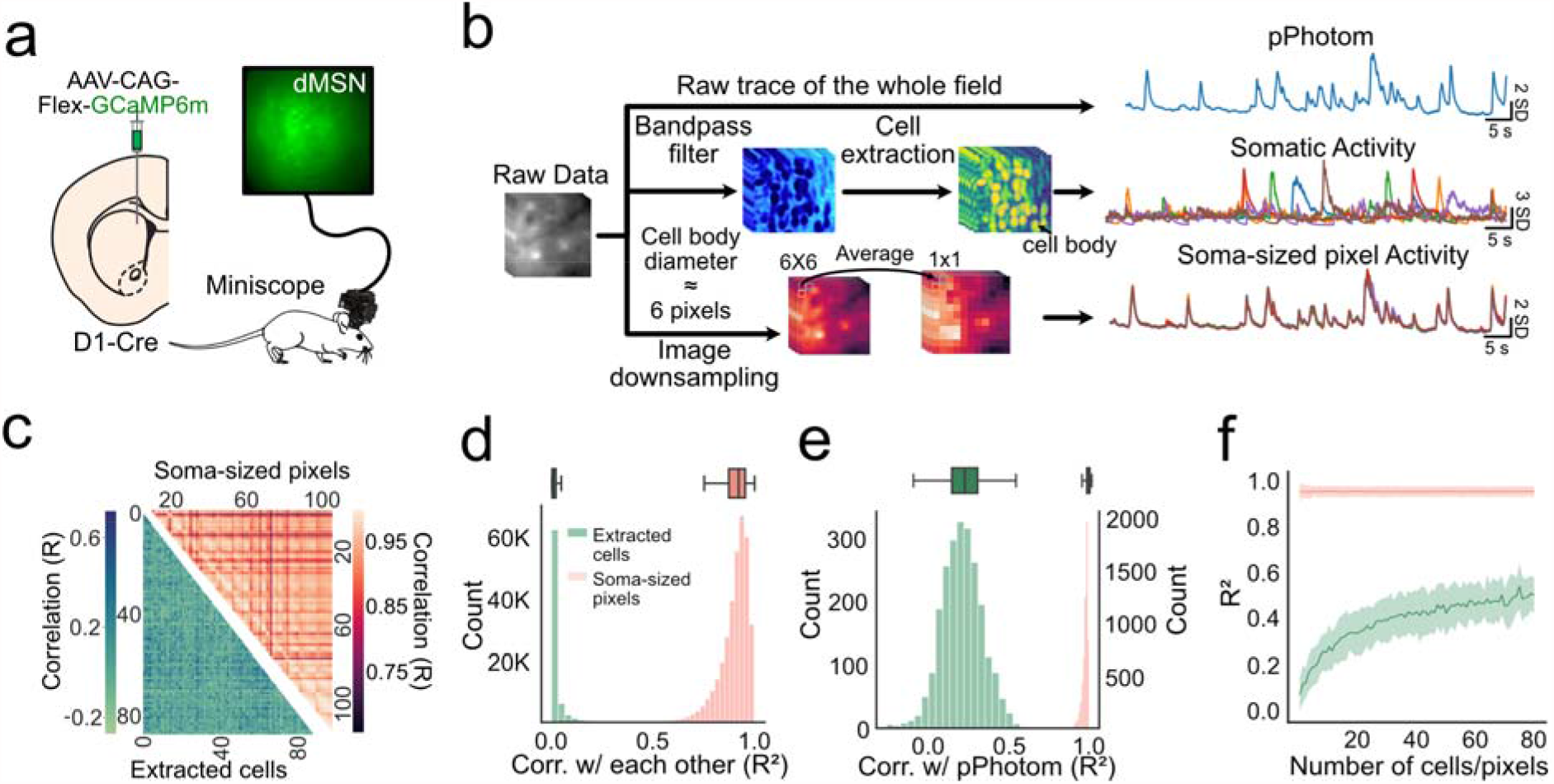
Fiber photometry correlates with whole-field changes in fluorescence signal. **a**, Experimental setup: D1-Cre mice were injected with Cre-dependent GCaMP6m in the DMS. **b**, Three signals were extracted from raw movies: 1) average of the entire field (pPhotom), 2) somatic signals (via CNMFe cell extraction), and 3) soma-sized regions (6×6 pixels) throughout the field. **c**, Heatmap showing the correlations among each extracted somatic signal (bottom), and among each soma-sized pixel (top). **d**, Quantification of **c. e**, Correlation between somatic signals and soma-sized pixels and pPhotom. **f**, same as **e**, but for increasing numbers of cells and pixels.

One potential confound of 1-photon endoscopic calcium imaging is that out of focus somas may not be detected by the extraction algorithm and may therefore contaminate the “non-somatic” signal. To eliminate this possibility, we performed volumetric 2-photon (2P) calcium imaging through a GRIN lens in dorsal striatum MSNs (n = 4 mice; **Fig. 3a**). Cell body locations (ROIs) were extracted from three different optical planes with the EXTRACT cell-extraction algorithm^13^. The contribution of out-of-focus somas to the average signal (pPhotom) was examined by comparing the correlations between pPhotom and masked movies that excluded somas from one, two, or all three optical planes (**Fig. 3b**). If out-of-focus somas contributed substantially to the pPhotom signal we would expect the correlation to be reduced as somas from different focal planes were excluded. This was not the case (one-way ANOVA, F = 1.12, p = 0.37; **Fig. 3c**).

**Figure 3.**
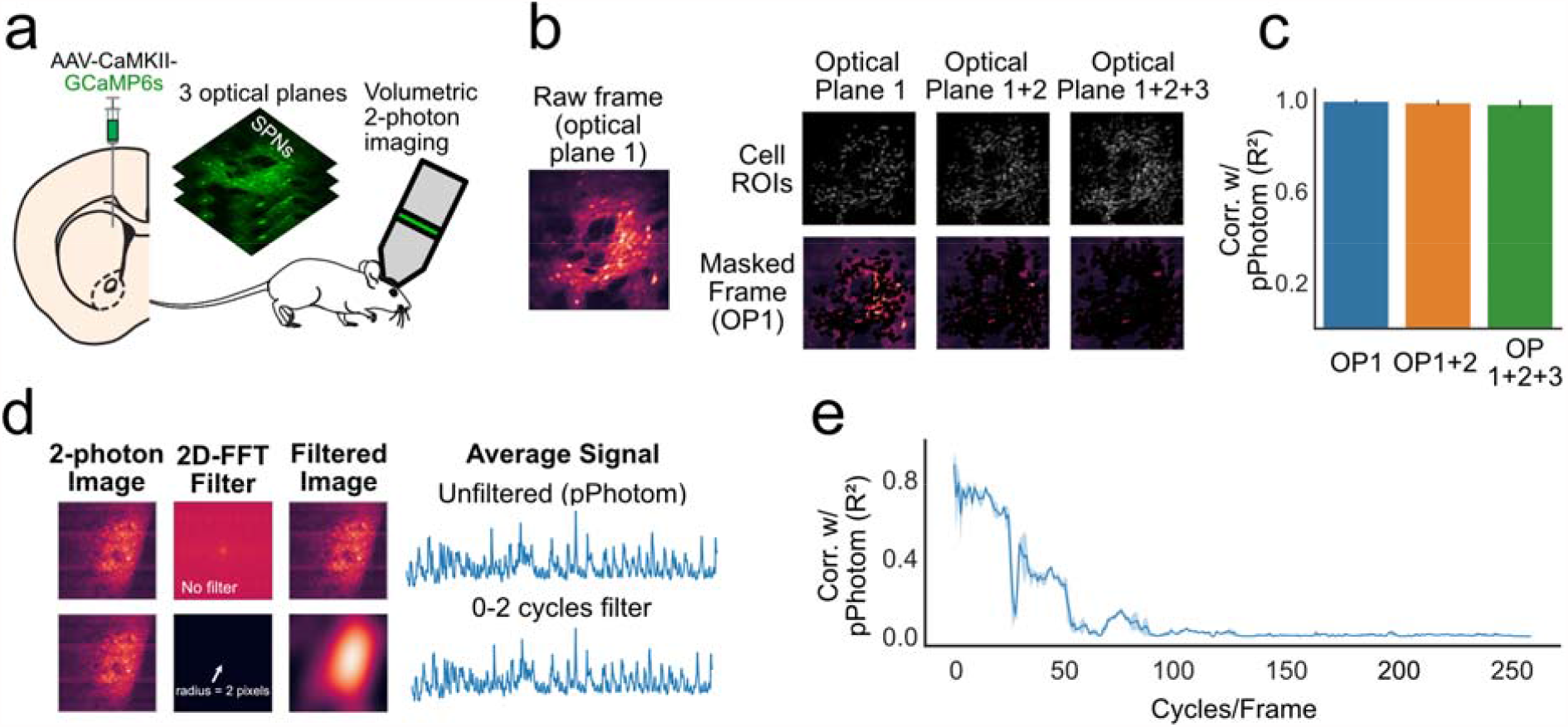
Out-of-focus cells do not contribute substantially to the fiber photometry signal. **a**, Experimental set-up: we expressed GCaMP6s in the DMS and performed volumetric two-photon imaging of three consecutive optical planes. **b**, The raw movie from optical plane 1 (OP1) was masked with somatic ROIs from either optical plane 1 only (OP1), or optical plane 1 and optical plane 2 (OP1+2), or from the three optical planes (OP1+2+3). **c**, Correlations between the average signal of the raw movie (pPhotom) and the masked movies **d**, 2D-FFTs were used to test the contribution of different spatial frequencies. Top row shows an example of the transformation between the time and space domain without applying any bandpass filter. Bottom row shows the same process but applying a bandpass filter that includes only the signal that is between 0 and 2 cycles per frame (full-frame). **e**, Correlations between the pPhotom signal and signal from different spatial frequencies (bin-width = 2 cycles/frame)

Finally, we applied spatial filters of different frequencies using 2D Fast Fourier Transformations (2D-FFTs) and tested how the resulting filtered movies correlated with the pPhotom signal (**Fig. 3d**). The pPhotom signal correlated best with low spatial frequencies, consistent with full-frame changes (*i*.*e*., at 0-2 cycles/frame, R^2^ > 0.85). Soma-sized spatial frequencies (ranging from 25-50 cycles/frame) correlated more weakly, at similar levels to average somatic signals in above analyses (**Fig. 3e**, R^2^ ∼ 0.4).

We conclude that the photometry signal from cell bodies in the striatum correlates more strongly with non-somatic than somatic calcium. This is consistent with recordings from cholinergic neurons in the striatum^10^ and may explain differences in the time-course of striatal photometry and spiking activity preceding actions^6^. This may be also be relevant to recent experiments that decoded position and speed information from hippocampal or cortical neuropil^15,16^. There are limitations to our study. First, the results presented here are limited to recordings from the striatum. Striatal neurons have extensive dendritic arbors that may accentuate the neuropil contribution to the fiber photometry signal^11,12^. Second, our photometry recordings were performed with GCaMP6s or GCaMP6m, which both have relatively slow kinetics^10^. Different relationships may be observed with GCaMP variants with faster kinetics, target GCaMP to specific cellular compartments^17,18^, or experiments that position the recording probe in axonal projection fields^19^. Despite these limitations, we argue that fiber photometry of cell bodies should not be interpreted as a proxy for spiking activity in a structure, but rather as primarily reflecting changes in neuropil calcium, which may therefore reflect inputs to a structure more so than outputs from that structure.

## Acknowledgements

We thank the HHMI GENIE project for GCaMP reagents and Meaghan Creed and Bridget Matikainen-Ankney for critical reading of the manuscript. Research supported by the Washington University Diabetes Research Center (DK020579), Nutrition Obesity Research Center (DK056341) and McDonnell Centers for Systems and Cellular Neuroscience.

## Author Contributions

**Legaria AA:** Conceptualization, Methodology, Software, Formal analysis, Investigation, Data Curation, Writing - Original Draft, Writing - Review & Editing, Visualization. **Yang B**: Data Curation, Formal analysis. **Ahanonu B**: Software, Methodology, Writing - Review & Editing. **Licholai JA**: Conceptualization, Methodology. **Parker JG**: Methodology, Data Curation, Writing - Review & Editing, Resources, Supervision. **Kravitz AV**: Conceptualization, Investigation, Writing - Original Draft, Resources, Supervision.

## Methods

### Subjects

The animals used in this study were 12 wildtype C57BL6 mice, 8 Drd1-Cre, 6 A2a-Cre, and 4 Drd1-Cre; Ai14 mice on a C57BL6/J background. Animals were housed in either the Washington University in St Louis animal facilities in standard vivarium cages with *ad libitum* food and water and a non-reversed 12-hour dark/light cycle or the Northwestern University animal facility with a reversed 12-hour light/dark cycle. All experimental procedures were approved by the Washington University Animal Care and Use Committee and the Northwestern University Animal Care and Use Committee.

### Viral transduction

Anesthesia was induced with 3-5% isoflurane and maintained at 0.5-1.5% isoflurane during stereotactic surgery. Ear bars and mouth holder were used to keep the mouse head in place while the skin was shaved and disinfected with a povidone/iodine solution. The skull was exposed and 1-mm diameter craniotomy was made with a microdrill mounted to the stereotaxic manipulator. Injections were performed with a glass pipette mounted in a Nanoject 3 infusion system (Drummond Scientific). 500 nL of virus AAV1-Syn-GCaMP6s AAV2/9-CAG-FLEX-GCaMP6m-WPRE.SV40 (1.37 × 10^12^ genome copies (GC) · ml^-1^ ; Penn Vector Core), or AAV2/9-CaMKII-GCaMP6s (1.2 × 10^12^ GC · ml^-1^) virus was infused over 10 minutes into either the dorsal striatum (AP +0.5 mm, ML +1.5 mm, DV -2.8 mm) or ventral striatum (AP +0.5 mm, ML +1.2 mm, DV -4.5 mm). The injector was left in place for 5 or 10 minutes before removal.

### Optical guide implantation and head bar placement

We used a 1.4-mm-diameter drill bit to create another craniotomy (AP=1.0mm; ML=1.5mm) for implantation of the optical guide tube. We fabricated this guide tube by using ultraviolet liquid adhesive (Norland #81) to fix a 2.5-mm-diameter disc of #0 glass to the tip of a 3.8-mm-long, extra-thin 18-gauge stainless steel tube (McMaster-Carr). We ground off any excess glass using a polishing wheel (Ultratec). Using a 27-gauge blunt-end needle, we aspirated the cortex down to DV=-2.1mm from the dura and implanted the exterior glass face of the optical guide tube at DV=-2.35mm. After stereotaxic placement of these components, we attached a headbar to the entire assembly using Metabond (Parkell) and dental acrylic. For mice used for two-photon imaging, we used additional dental acrylic to construct a reservoir for holding water for the water-immersion objective lens. Mice recovered for 3–4 weeks before two-photon imaging experiments or mounting of the miniature microscope.

### Gradient Index (GRIN) lens implantation and mounting of miniature microscope

After 3-4 weeks, we inserted a gradient refractive index (GRIN) lens (1mm diameter; 4.12mm length; 0.46 numerical aperture; 0.45 pitch; GRINTECH GmbH or Inscopix Inc.) into the optical guide tube. In mice with uniform indicator expression, we secured the GRIN lens in the guide tube with ultraviolet (UV)-light curable epoxy (Loctite 4305). For miniscope imaging, after affixing the GRIN lens, we lowered a miniature microscope (nVistaHD, Inscopix Inc.) towards the GRIN lens until the fluorescent tissue was in focus. To secure the miniature microscope to the cranium, we created a base on the cranium around the GRIN lens using blue-light curable resin (Flow-It ALC; Pentron). We attached the base plate of the miniature microscope to the resin base using UV-light curable epoxy (Loctite 4305). After affixing its base plate, we released the microscope and attached a base plate cover (Inscopix Inc.). We coated the resin with black nail polish (Black Onyx, OPI) to make it opaque.

### Implantation of electrode arrays

Following viral infusion, a combined electrophysiology/fiber photometry device was implanted. Fiber optic cannulae (200 μm, 0.50 NA) with 1.25mm ceramic ferrules were purchased from Thorlabs and cut to 6mm long. These cannulae were mounted in a custom electrode array with 32 Teflon-coated tungsten microwires (35 µm diameter; Innovative Neurophysiology) that positioned the wires in a semi-circle surrounding a central gap where the photometry fiber was mounted. This combined photometry/electrical recording device was implanted into the right DMS (AP, +0.5 mm, ML, +1.5 mm, DV, -2.8 mm). The device was secured to skull with a thin layer of adhesive dental cement (C &B Metabond, Parkell) followed by a larger layer of acrylic dental dement (Lang Dental). After the cement fully cured animals were placed back in their home-cage on a pre-heated pad at 37°C. After recovery, animals received a subcutaneous injection of meloxicam (10 mg · kg^-1^) and were returned to their home-cages for recovery. Mice recovered for at least 2 weeks to allow for viral expression prior to recording.

### Electrophysiological recordings

Neurophysiological signals were recorded by a multi-channel neurophysiology system (Plexon Omniplex, Plexon Inc). Spike channels were acquired at 40 kHz and band-pass filtered at 150 Hz to 3 kHz before spike sorting. Recordings were performed in a 9” x 12” clear plastic box and lasted between one and three hours. Video and tracking data was also recorded in real-time with the Plexon Cineplex system.

### Fiber photometry recordings

Fiber photometry acquisition was performed with a Neurophotometrics fiber photometry system (FP3001, Neurphotometrics LTD). Briefly, this system utilizes a 470nm blue light LED which was left on continuously at 40-100uW to excite GCaMP, and a fluorescence light path that includes a dichroic mirror to pass emitted green fluorescence to a CMOS camera (FLIR BlackFly). Fluorescence signals from the camera are processed with Bonsai (https://bonsai-rx.org/docs) and transmitted as a voltage signal to the Plexon Omniplex for simultaneous digitizing with the electrophysiological data.

### Electrophysiology/Fiber photometry data analysis

Single and multiunits were manually discriminated using principal component analysis (Offline Sorter; Plexon), using MANCOVA analyses to determine if single unit clusters were statistically distinct from multi-unit clusters. Where single unit isolation did not reach statistical significance, spike clusters were combined into multiunits. Data analysis was performed using a custom Python/Neuroexplorer pipeline (code available at: https://osf.io/8j7g2/), described below.

#### Photometry signal preprocessing

The fiber photometry signal was processed using a custom python/Neuroexplorer5 pipeline (code available at: https://osf.io/8j7g2/). We first applied both a lowpass (6Hz) and a highpass filter (0.0005Hz) to the photometry signal to correct for high frequency noise and photo bleaching, respectively. Photometry peaks were then identified using the scipy.signal.find_peaks function, setting a prominence threshold of 1. Photometry peak amplitude quantiles were created by diving the peak timestamps into five groups containing an equal number of peaks each, using the pandas function pd.qcut.

#### Burst Analysis

Only recordings that had at least 10 units were used for the burst analysis. From these recordings, 10 units were pseudo-randomly (avoiding very high/low firing rate) picked and their firing activity was pooled. Bursts of spiking were detected using the Burst Analysis tool in Neuroexplorer, using a firing rate based algorithm. The parameters were set as follows: Hisotgram Bin: 0.05 seconds; Guassian Filter width: 5 bins; Burst Detection Threshold: 3 standard deviations; Minimum number of spikes in burst: 5.

#### Normalization of signals

In all peri-event histograms (PEHs), all the signals were z-scored to a baseline period encompassing the first 25% of the analyzed interval.

#### Classification of responsive units

The classification of responsive units in Figure 2 F-G was done by creating a PEH of each neuron around the photometry peak (figure 2D). We set a baseline period for each neuron (−4 to -2 seconds from the photometry peak) and considered a neuron to be responsive if firing in a signal window (−1 to 1 seconds from the photometry peak) was at least 3 z-scores higher than the baseline window for at least 250 consecutive milliseconds. We chose 3 standard deviations based on a permutation analysis that detected <1% false positives with this threshold (data not shown).

### One-photon endoscopic calcium imaging recordings

Brain imaging in freely moving mice occurred in a circular arena (31cm in diameter). To habituate mice to this arena, mice explored it for 1h on each of three sequential days before any Ca^2+^ imaging. Prior to each imaging session we head-fixed each mouse by its implanted head bar and allowed the mouse to walk or run-in place on a running wheel. We then attached the miniature microscope and adjusted the focal setting to optimize the field-of-view. After securing the microscope to the head of the mouse, we detached the mouse from its head restraint and allowed it to freely explore the circular arena. After allowing ≥10min for the mouse to habituate to the arena, fluorescence Ca^2+^ imaging commenced using 50–200μW of illumination power at the specimen plane and a 20-Hz frame acquisition rate.

### Endoscopic calcium imaging analysis

#### Cropping

To better match the surface area of the most commonly used fiber photometry fiber, we cropped each 1mm GRIN lens images into six 200micron square regions with the FIJI distribution of ImageJ^1^.

#### Somatic activity extraction

We used the CaImAn^2^ cell body extraction Jupyter notebook pipeline to extract somatic activity from 1P movies. Briefly, this pipeline implements motion correction and the CNMF-E algorithm^3^ in an online notebook, returning quality metrics and images of extracted somatic signals for subsequent analysis. We then averaged the activity trace of somatic signals.

#### Non-somatic activity extraction

After the ROIs of cell bodies were extracting using the CaImAn pipeline, these ROIs were used to mask activity coming from those ROIs. The remaining activity was averaged for each frame of the movie to create an average non-somatic signal.

#### Extraction of photometry signals

The proxy for photometry signal was obtained by averaging the intensity of the entire field for each frame of the raw movie.

### Two-photon calcium imaging recordings

Drd1-Cre; Ai14 mice injected with AAV2/9-CaMKII-GCaMP6s were used for two-photon calcium imaging. GCaMP6s was constitutively expressed in both MSN types, whereas tdTomato expression was restricted to D1-MSNs. After habituating mice to head-fixation on a running wheel, we used two-photon microscope with a piezoelectric actuator (Bruker) to acquire movies of Ca^2+^ activity and tdTomato expression at three imaging planes separated by 20 µm in the dorsal striatum of head-fixed mice during wheel running. We used a tunable laser (Insight X3, Spectra Physics) and a 16x/0.8NA objective (Nikon) to acquire 512 × 512 pixel movies of each plane at a 30-Hz frame acquisition rate (effectively 6 Hz per plane). We used λ = 920 nm to simultaneously excite tdTomato and GCaMP6s fluorescence which we detected using GaAsP photomultiplier tubes and band-pass filters (520/40 for GCaMP6s and 595/50 for tdTomato).

### Two-photon calcium imaging processing

We used an exponential fit to normalize slow variations in green and red fluorescence intensity assumed to be due to photobleaching. We then motion corrected the tdTomato movie using NormCorre^4^. We then applied the tdTomato motion correction transformations to the movie frames of the green GCaMP6s fluorescence movie. We then corrected for fluctuations in background fluorescence intensity in the GCaMP6s movie by applying a Gaussian low-pass filter to each image, then dividing each image frame by its low-pass filtered version. We then down sampled the GCaMP6s movie by a factor of 2 via linear interpolation.

### Two-photon calcium imaging analysis

Data analysis was performed using a custom Python/Neuroexplorer pipeline (code available at: https://osf.io/8j7g2/), described below:

#### Fast Fourier Transformations (FFTs)

All FFTs analyses were done using the numpy.fft2 module of numpy. To obtain the signal coming from different spatial frequencies, an FFT was applied to each frame, then a bandpass filter was applied through a circular mask that selected for specific spatial frequencies. Finally, an inverse FFT was performed to recreate a filtered image that contained only the spatial frequencies allowed by the filter.

#### Non-somatic activity extraction

After the ROIs of cell bodies were extracting using the EXTRACT pipeline, these ROIs were used in the raw movie to mask all the activity coming from those ROIs. The remaining activity was then averaged for each frame of the movie.

### Histology

At the end of the experiments, we performed histological verification of implant placements. Animals were anesthetized with isoflurane, decapitated, and their brains were quickly removed and placed in 10% formalin solution. Brains were incubated overnight in 10% formalin solution, and then moved to 30% sucrose solution until sectioning. Coronal slices containing the striatum were prepared using a freezing microtome (Leica SM2010R). Slices were mounted on microscope slides with a mounting media and imaged with an epifluorescence microscope (Zeiss). For in vivo electrophysiology electrode placement was assessed via observation of implant tract or electric-lesions that were made under anesthesia before decapitation (performed with a 5-s long pulse of 10 mA; Ugo Basile Lesion Making Device).

